# Tumor-free elongation of mammalian nephrogenesis by excess fetal GDNF

**DOI:** 10.1101/2020.05.28.120865

**Authors:** Hao Li, Jussi Kupari, Yujuan Gui, Edward Siefker, Benson Lu, Kärt Mätlik, Soophie Olfat, Ana R Montaño-Rodríguez, Sung-Ho Huh, Franklin Costantini, Jaan-Olle Andressoo, Satu Kuure

**Affiliations:** Stem Cells and Metabolism Research Program, Faculty of Medicine, University of Helsinki, Finland; Institute of Biotechnology, Helsinki Institute of Life Science, University of Helsinki, Finland; Division of Molecular Neurobiology, Department of Medical Biochemistry and Biophysics, Karolinska Institutet, SE-17177, Sweden; Department of Life Sciences and Medicine, University of Luxembourg, L-4367, Luxembourg; Department of Developmental Neuroscience, University of Nebraska Medical Center, NE, USA; Department of Genetics and Development, Columbia University Medical Center, NY, USA; Department of Pharmacology, Faculty of Medicine, University of Helsinki, Finland; Department of Neurobiology, Care Sciences and Society, Karolinska Institutet, Sweden; GM-unit, Laboratory Animal Centre, Helsinki Institute of Life Science, University of Helsinki, Finland

**Author notes:** Co-last author (Jaan-Olle Andressoo, Satu Kuure).

## Abstract

Due to poor regenerative capacity of adult kidneys, nephron endowment defined by the nephrogenic program during the fetal period dictates renal and related cardiovascular health throughout life. We show that the neurotropic factor GDNF, which is in clinical trials for Parkinson’s disease, is capable of prolonging the nephrogenic program beyond its normal cessation without increasing the risk of kidney tumors. Our data demonstrates that excess GDNF expands the nephrogenic program by maintaining nephron progenitors and nephrogenesis in postnatal mouse kidneys. GDNF, through its transcriptional targets excreted from the adjacent epithelium, has a major effect on nephron progenitor self-renewal and maintenance; abnormally high GDNF inhibits nephron progenitor proliferation, but lowering its level normalizes the nephrogenic program to that permissive for nephron progenitor self-renewal and differentiation. Based on our results, we propose that the lifespan of nephron progenitors is determined by mechanisms related to perception of GDNF and other signaling levels.

## INTRODUCTION

The mammalian kidney is a non-regenerating, blood filtrating organ with essential detoxification and electrolyte balancing functions. The nephrons executing the main renal functions are all generated during the fetal period, while adult kidneys show only limited repair capacity and compensatory growth through enlargement of glomeruli (hypertrophy) (Chawla & Kimmel, 2012, Humphreys, Valerius et al., 2008, Rinkevich, Montoro et al., 2014, Romagnani, Lasagni et al., 2013). Reduction in nephron mass is a widely accepted risk factor for hypertension, proteinuria, glomerulosclerosis and end-stage renal disease (Barker, Shiell et al., 2000, Bertram, Douglas-Denton et al., 2011, Hoy, Bertram et al., 2008, Keller, Zimmer et al., 2003, Lackland, Bendall et al., 2000, Little & Bertram, 2009, Schreuder, 2012, Schreuder, Langemeijer et al., 2008). Thus, congenital nephron endowment greatly influences renal health throughout the individual’s lifespan.

The nephrogenic program ends at gestation week 36 in humans and postnatal day (P3) in mouse (Cebrian, Asai et al., 2014, Cullen-McEwen, 2016, Hartman, Lai et al., 2007, Hinchliffe, Sargent et al., 1991, Lindstrom, McMahon et al., 2018, Rumballe, Georgas et al., 2011, Short, Combes et al., 2014), but the mechanisms contributing to the cessation of nephrogenesis are poorly understood. It has been postulated that stage-specific changes in growth patterns (Cebrian, Borodo et al., 2004) and cell intrinsic age-sensing mechanisms (Chen, Brunskill et al., 2015a) are involved. Molecularly, bone morphogenetic protein (BMP)-induced SMAD signaling (Brown, Muthukrishnan et al., 2015), hamartin (*Tsc1*) (Volovelsky, Nguyen et al., 2018) and microRNA regulation (Yermalovich, Osborne et al., 2019) participate in defining the duration of the nephrogenic program. However, plasticity of the nephrogenic program has not been fully explored.

Nephrons differentiate from an unipotent nephron progenitor (NP) pool also known as metanephric or cap mesenchyme (CM). NPs are present in the embryonic kidney where they are found around each ureteric bud (UB) tip since its establishment. NP cells express a unique repertoire of markers including SIX2, glial cell line derived neurotrophic factor (GDNF) and CITED1 (Boyle, Misfeldt et al., 2008, Cebrian et al., 2014, Kobayashi, Valerius et al., 2008, Self, Lagutin et al., 2006), while the adjacent mesenchyme is mainly populated by the interstitial/stromal progenitors expressing e.g. MEIS1/2 and FOXD1 (Hatini, Huh et al., 1996, Humphreys, Lin et al., 2010). NPs represent a self-renewing, multipotent progenitor population, which is capable of giving rise to all epithelial components of the mature nephron (Kobayashi et al., 2008, Lindström, Guo et al., 2018, Little, 2016, Mugford, Yu et al., 2009, Takasato & Little, 2015). Nephrogenesis is synchronized with the UB branching, which simultaneously maintains the undifferentiated NP population and induces a subset of NPs to differentiate (O’Brien, 2018). This occurs through well-defined precursor stages covering the initial condensation, formation of pretubular aggregates (PA), and mesenchyme-to-epithelium transition (MET) into renal vesicles (RV) and comma- and S-shaped bodies (CSB, SSB), which finally mature into all nephron segments (Saxen, 1987).

The balance in self-renewal versus differentiation within the NP population depends on signaling executed by the classical reciprocal inductive tissue interactions between the UB and the neighboring mesenchyme. The signals originating from the NPs, such as GDNF and fibroblast growth factors (FGFs), initiate the UB morphogenesis, through which the embryonic kidney grows in size, achieves its shape and establishes the collecting duct system (Costantini & Kopan, 2010, Hendry, Rumballe et al., 2011). GDNF-activated RET signaling in the UB tips regulates a transcriptional profile that is responsible for the control of collecting ductal progenitors and their differentiation morphogenesis (Kurtzeborn, Cebrian et al., 2018, Li, Jakobson et al., 2019, Lu, Cebrian et al., 2009, Ola, Jakobson et al., 2011, Riccio, Cebrian et al., 2016). GDNF regulated genes also comprise secreted proteins, such as WNT11 and CRLF1 with demonstrated potential to reciprocally regulate the nephrogenic program (O’Brien, Combes et al., 2018, Schmidt-Ott, Yang et al., 2005). Other important UB-derived nephrogenesis regulators are WNT9b and FGF-ligands, which influence NP expansion and differentiation (Barak, Huh et al., 2012, Carroll, Park et al., 2005, Karner, Das et al., 2011, Kiefer, Robbins et al., 2012).

We previously reported that the increase in endogenous expression of GDNF at the post-transcriptional level results in renal hypoplasia (Kumar, Kopra et al., 2015, Li et al., 2019). Unlike the *Gdnf* inactivation model, the *Gdnf* ^hyper/hyper^ mice do not die shortly after birth but survive up to three weeks of age (Kumar et al., 2015, Li et al., 2019, Moore, Klein et al., 1996, Pichel, Shen et al., 1996). Here we demonstrate that excess GDNF expands the nephrogenic program beyond its normal cessation by maintaining NP cells and nephrogenesis without neoplastic growth in postnatal mouse kidneys, unlike as reported for microRNA *let-7* and RNA-binding protein *Lin28* modulation (Urbach, Yermalovich et al., 2014, Yermalovich et al., 2019). Our results approve remarkable plasticity in mammalian kidneys, which could possibly be utilized in regenerative medicine in the future.

## RESULTS

### The embryonic nephrogenic program depends on GDNF

The spatial arrangement of SIX2-positive NPs in the embryonic and early postnatal kidneys follows a stereotypic, well described pattern, where the initial multi-cell-layered tissue thins over time without much affecting the overall niche organization (Lindstrom et al., 2018, Short et al., 2014). Quantification of the SIX2-positive NP cell numbers at the on-set of renal development revealed an increase in the initial progenitor pool (Supplementary figure 1A-C). At the initiation of UB branching, NP amount in *Gdnf*^hyper/hyper^ kidneys resembled that in wild type kidneys (Figure 1A-B, Supplementary figure 1D) but were quickly decreased due to 57% reduced NP cell mitosis in the presence of endogenous (Supplementary figure 1E) and exogenous excess GDNF (Supplementary figure 1F). Accordingly, *Gdnf*^hyper/hyper^ kidneys soon exhibited a thinned and discontinuous layer of SIX2-positive NPs in both culture and *in vivo* models (Figure 1 C-F). PAX2 staining, labeling not only NPs but also the early nephron precursors, demonstrated similar dramatic reduction in NPs and additional gradual decrease in differentiating nephrons (Supplementary figure 2). Collectively, the data show that excess GDNF, which primarily functions in the UB due to its receptor complex localization (Pachnis, Mankoo et al., 1993), induces a rapid drop-off in NP cell proliferation.

**Figure 1.**
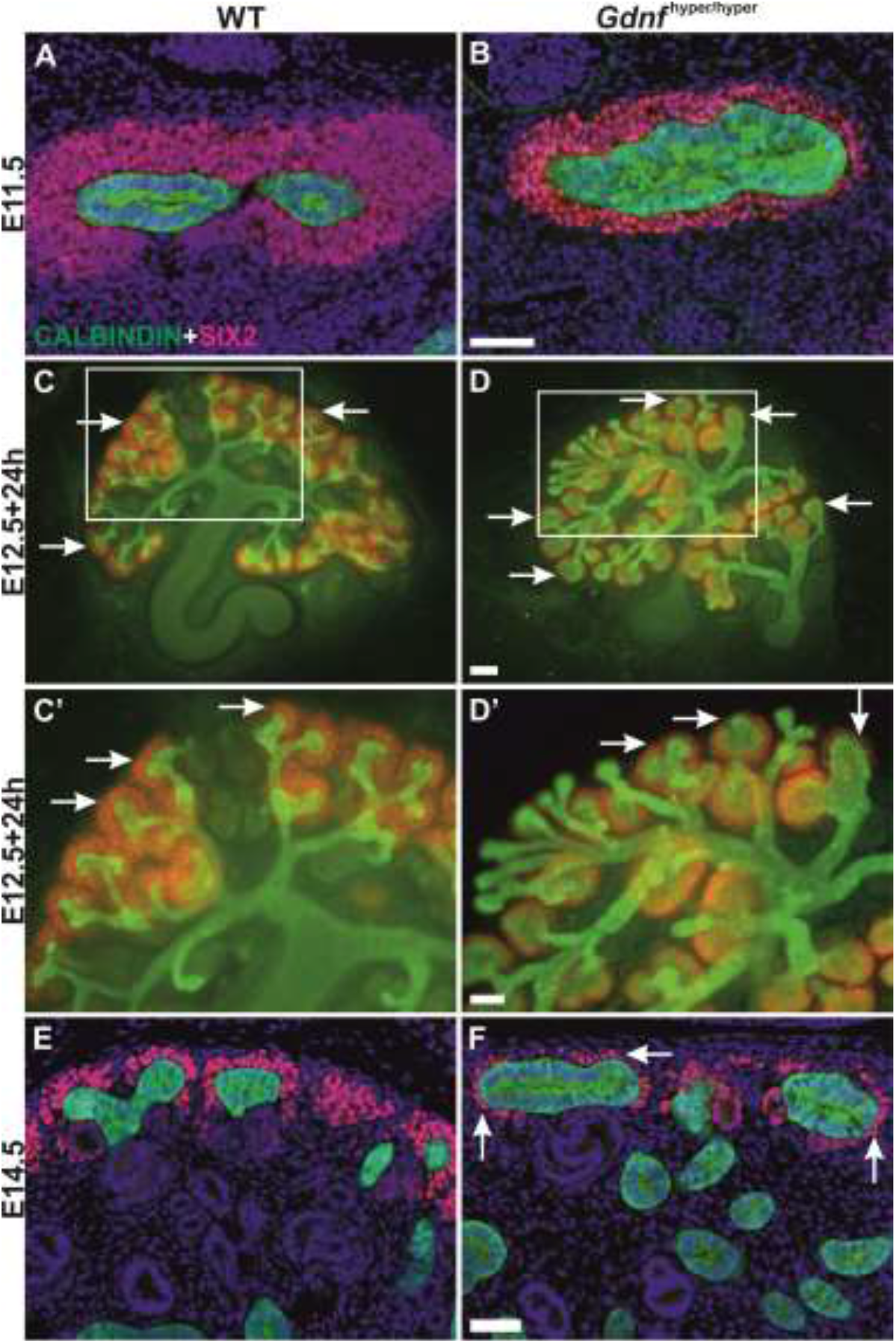
Nephron progenitor depletion in embryonic *Gdnf* ^hyper/hyper^ kidneys. (A) Nephron progenitor marker SIX2 (red) localizes to mesenchyme capping ureteric bud tips (CALBINDIN, green) in WT and (B) *Gdnf*^hyper/hyper^ kidneys at E11.5. (C) E12.5 WT and (D) *Gdnf*^hyper/hyper^ kidneys cultured *in vitro* for 24h and stained for SIX2 (red) to visualize nephron progenitor population and CALBINDIN (green) for ureteric bud. C’ and D’ show zoom in to the cortical kidney region where the tips and progenitor population are located. Nephron progenitors are abundant in WT E14.5 kidney while clearly reduced (arrows) in (F) *Gdnf*^hyper/hyper^ kidney of the same stage. Arrows point to the nephron progenitor cells (red). *Scale bar*: 100μm.

In addition to the impairment in NP self-renewal and propagation, prematurely accelerated NP differentiation could contribute to the decline in NP proportion in their niche. To assess this, nephrogenesis was examined by staining with LEF1, which labels the earliest differentiation-committed cells, and with CYCLIN D1 labeling the nephrons undergoing full epithelialization (Lindstrom, Carragher et al., 2015). This revealed a general decrease in early nephron precursors and indicated hysteresis in the subsequent differentiation of *Gdnf*^hyper/hyper^ nephrons (Figure 2). Together these data indicate that in fetal kidneys, excess GDNF negatively regulates embryonic NP cell self-renewal and consequently diminishes the number of differentiating nephron precursors.

**Figure 2.**
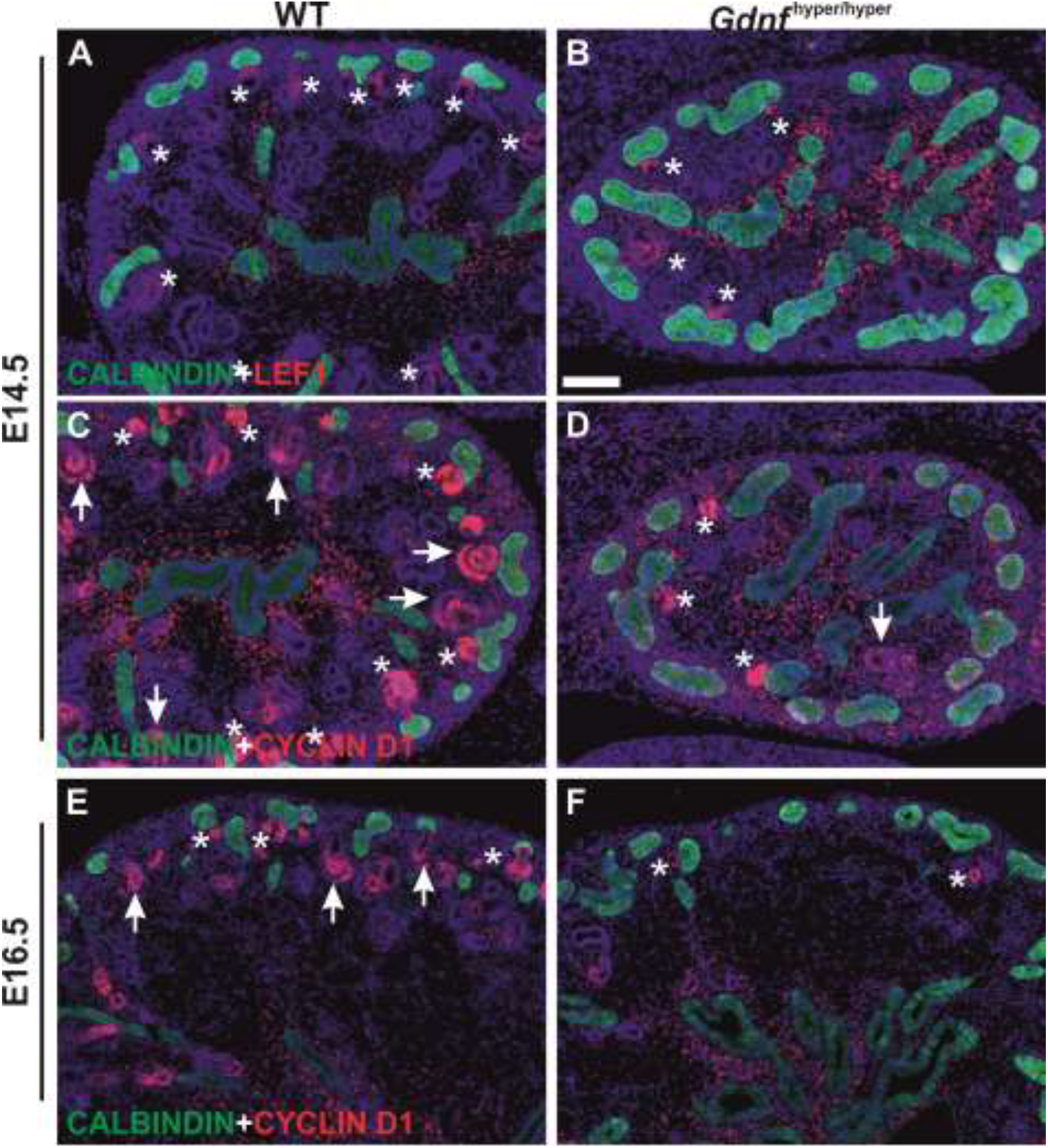
Effect of excess GDNF on embryonic nephrogenesis. (A) LEF1 (red), a marker of pretubular aggregates, distal renal vesicles and S-shaped bodies, in WT E14.5 kidney. (B) E14.5 *Gdnf*^hyper/hyper^ kidney shows very few LEF1-positive differentiating nephron precursors and typical abundance of ureteric bud epithelium as detected by CALBINDIN staining (green). (C) CYCLIN D1 (red) stains all precursors of differentiating nephrons (asterisk marks pretubular aggregates and comma-shaped bodies, arrow points to S-shaped bodies), which can be observed abundantly next to ureteric bud epithelium (green) in E14.5 WT kidney. (D) Significantly fewer differentiating nephron precursors are detected in E14.5 *Gdnf*^hyper/hyper^ kidney. (E) CYCLIN D1 (red) localization in E16.5 WT and (F) *Gdnf*^hyper/hyper^ kidneys. CALBINDIN (green) staining visualizes ureteric bud epithelium in all images. *Scale bar*: 100μm.

### Excess GDNF prolongs the lifespan of the postnatal nephron progenitor pool

Many mouse models with deficiency in kidney growth, either due to UB morphogenesis and/or nephron differentiation defect, die immediately after birth (Brown et al., 2015, Carroll & Das, 2013, Costantini & Kopan, 2010, O’Brien & McMahon, 2014). Despite the significantly reduced kidney size (Kumar et al., 2015, Li et al., 2019) and severely disrupted embryonic nephrogenesis (Figure 2), *Gdnf*^hyper/hyper^ mice survive up to three weeks after birth providing a possibility to examine postnatal nephrogenesis in this model.

In agreement with several previous studies (Brown et al., 2015, Cebrian et al., 2014, Rumballe et al., 2011, Short et al., 2014, Togel, Valerius et al., 2017, Yermalovich et al., 2019), analysis of the NP populations at the final stages of the nephrogenic program (postnatal days 1-10, P1-10) showed that NPs were principally lost from the wild type kidneys by P3, and only remnants of SIX2-positivity was detected in the differentiating nephron precursors (Figure 3A). Interestingly, the SIX2-postive NPs in most NP niches of *Gdnf*^hyper/hyper^ kidneys outnumbered those in wild type kidneys at P3 (Figure 3A-B, N≥3). Similarly, PAX2-positive NPs still capped the UB in *Gdnf*^hyper/hyper^ kidneys at P3, while in wild type kidneys the PAX2-positive cells located exclusively in the differentiating nephron precursors (Figure 3C-D).

**Figure 3.**
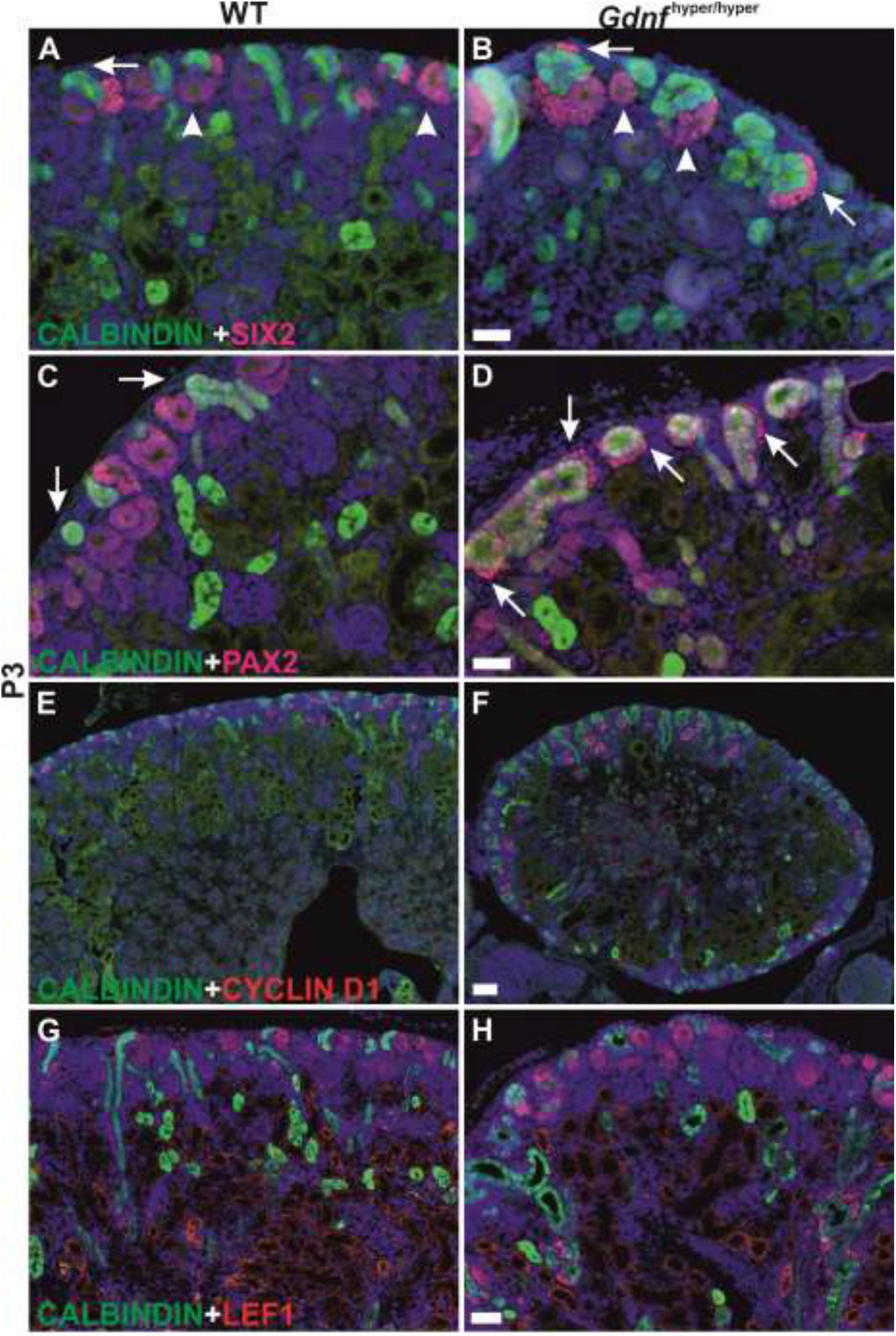
Nephron differentiation at the cessation of renal morphogenesis. (A) Postnatal day 3 (P3) WT kidney no longer hosts cortical nephron progenitors as shown by the loss of SIX2-positive cells in the cap mesenchyme (arrow). Instead, SIX2 localizes to lateral mesenchyme and early nephron precursors (arrowhead). (B) Cap mesenchyme positive for SIX2 is still present in *Gdnf*^hyper/hyper^ kidney at P3 (arrows). (C) PAX2 (red) and CALBINDIN (green) staining in WT and (D) *Gdnf*^hyper/hyper^ kidneys at P3. Arrows point to the position where progenitor cells are maintained in *Gdnf*^hyper/hyper^ kidneys. (E) Comparable amount of CYCLIN D1-positive (red) nephron precursors in P3 WT and (F) *Gdnf*^hyper/hyper^ kidneys. (G) Visualization of LEF1-positive nephron precursors (red) and ureteric epithelium (green) reveals similar ongoing nephrogenesis in WT and (H) *Gdnf*^hyper/hyper^ kidneys. *Scale bar*: 100μm.

Characterization of nephrogenesis in *Gdnf*^hyper/hyper^ kidneys at this stage demonstrated significant improvement from that in embryonic kidneys, as similar patterns of nephron precursors were detected both in wild type and *Gdnf*^hyper/hyper^ kidneys (Figure 3E-H; compare to Figure 2). Finally, we examined the presence of NP population at the later stages to find out that SIX2-positive NP cells were sustained until P6 in *Gdnf*^hyper/hyper^ kidneys (Figure 4, N≥3). This demonstrates that the NP existence period *in vivo* is not pre-fixed, and shows that modulation of growth factor levels, specifically GDNF, defines the timing of the final nephron differentiation wave.

**Figure 4.**
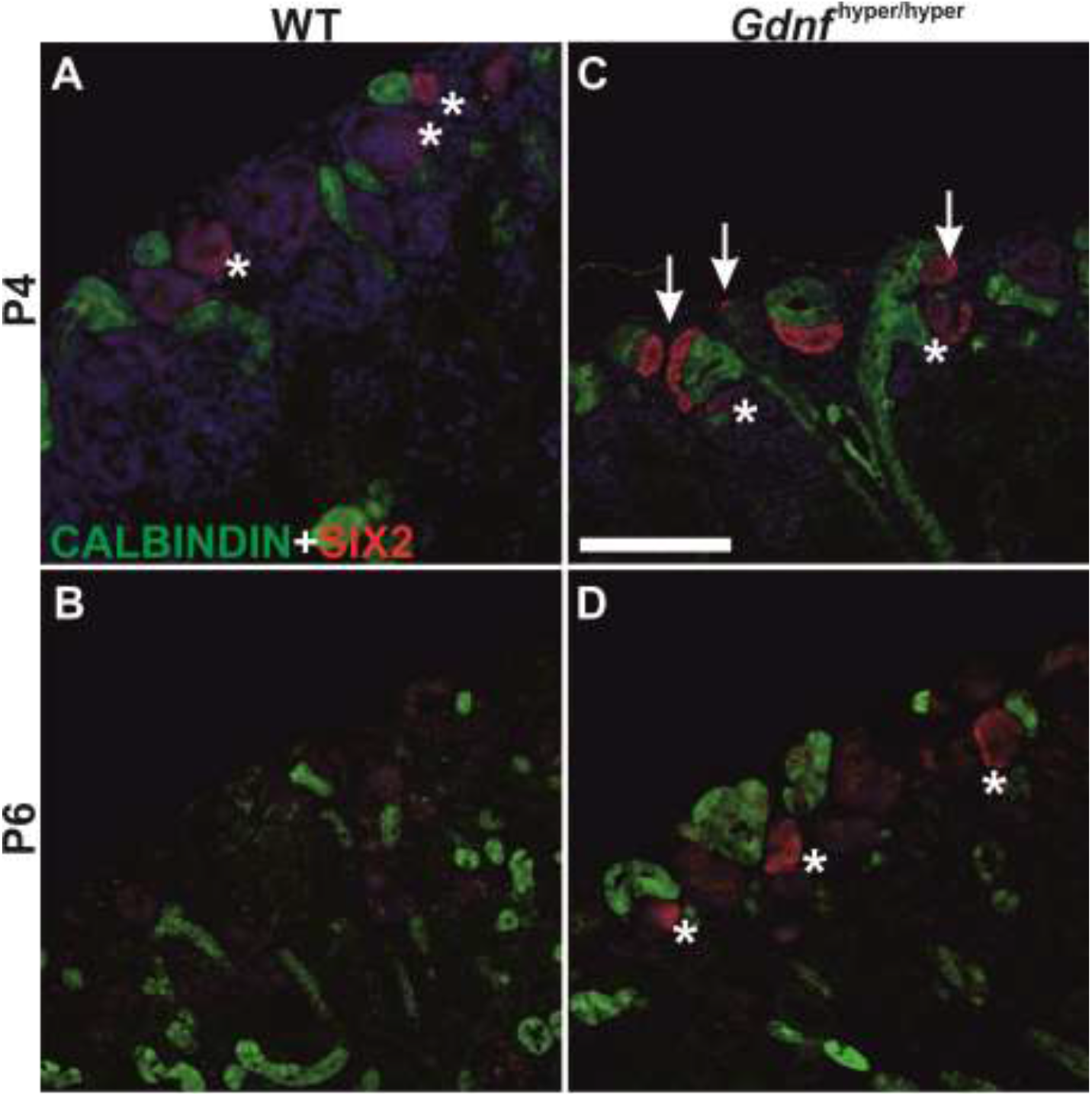
Nephron progenitors are persistently sustained in the postnatal *Gdnf*^hyper/hyper^ kidneys. (A) Localization of nephron progenitor marker SIX2 (red) in P4 WT and (B) *Gdnf*^hyper/hyper^ kidneys shows loss of progenitors in WT kidneys where SIX2 is found only in weakly positive renal vesicles (asterisk), while in *Gdnf*^hyper/hyper^ kidney it also localizes to mesenchymal progenitors capping the ureteric bud (green) tips (arrows). (C) Localization of SIX2 (red) and CALBINDIN (green) in P6 WT and (D) *Gdnf*^hyper/hyper^ kidneys reveals the persistence of nephron progenitor cells in the mutant kidney. *Scale bar:* 100μm.

We hypothesized that persistent NP cells in postnatal *Gdnf*^hyper/hyper^ kidneys could either be caused by the GDNF-specific effects on the NP population, or alternatively, result from general compensatory mechanisms provoked by renal hypoplasia caused by the ureteric bud branching defect(Kumar et al., 2015, Li et al., 2019). To discriminate between the two options, postnatal NPs in hypoplastic kidneys of *Fgf9;20* deficient mice were first examined. Those animals show similar thinning of embryonic NP cell layers (Barak et al., 2012) as detected in *Gdnf*^hyper/hyper^ kidneys. The analysis failed to detect SIX2- and/or PAX2-positive cells in the typical NP niche localization of *Fgf9;20* deficient kidneys (Supplementary figure 3). This, together with an extensive literature examination, which did not, to the best of our knowledge, disclose sustained NPs in other hypoplastic kidney models except for tumorigenic micro-RNA *Lin28/Let-7* modulation (Urbach et al., 2014, Yermalovich et al., 2019) supports the view that NPs are maintained in postnatal kidneys due to excess expression of GDNF.

### Sustained postnatal nephron progenitors maintain the nephrogenic program beyond its normal cessation

To assess the differentiation potential of NPs maintained in the postnatal *Gdnf*^hyper/hyper^ kidneys, nephrogenesis was studied by analyzing proliferation and nephron precursor abundance after its normal cessation. Ki67 staining revealed comparable postnatal proliferation in wild type and *Gdnf*^hyper/hyper^ kidneys until P4, despite the overall abnormal renal morphology in mutant pups (Figure 5A-B). From P5 onwards the Ki67-positive cells decreased dramatically in the cortical differentiation zone of wild type kidneys (Figure 5C, E and G). Interestingly, such a drop in overall proliferation was not detected in *Gdnf*^hyper/hyper^ kidneys, which showed actively cycling cells in nephron precursor-like structures still at P7 (Figure 5D, F and H). The higher cortical proliferation in *Gdnf*^hyper/hyper^ kidneys was no longer detected at postnatal day 12 (Supplementary figure 4A-B).

**Figure 5.**
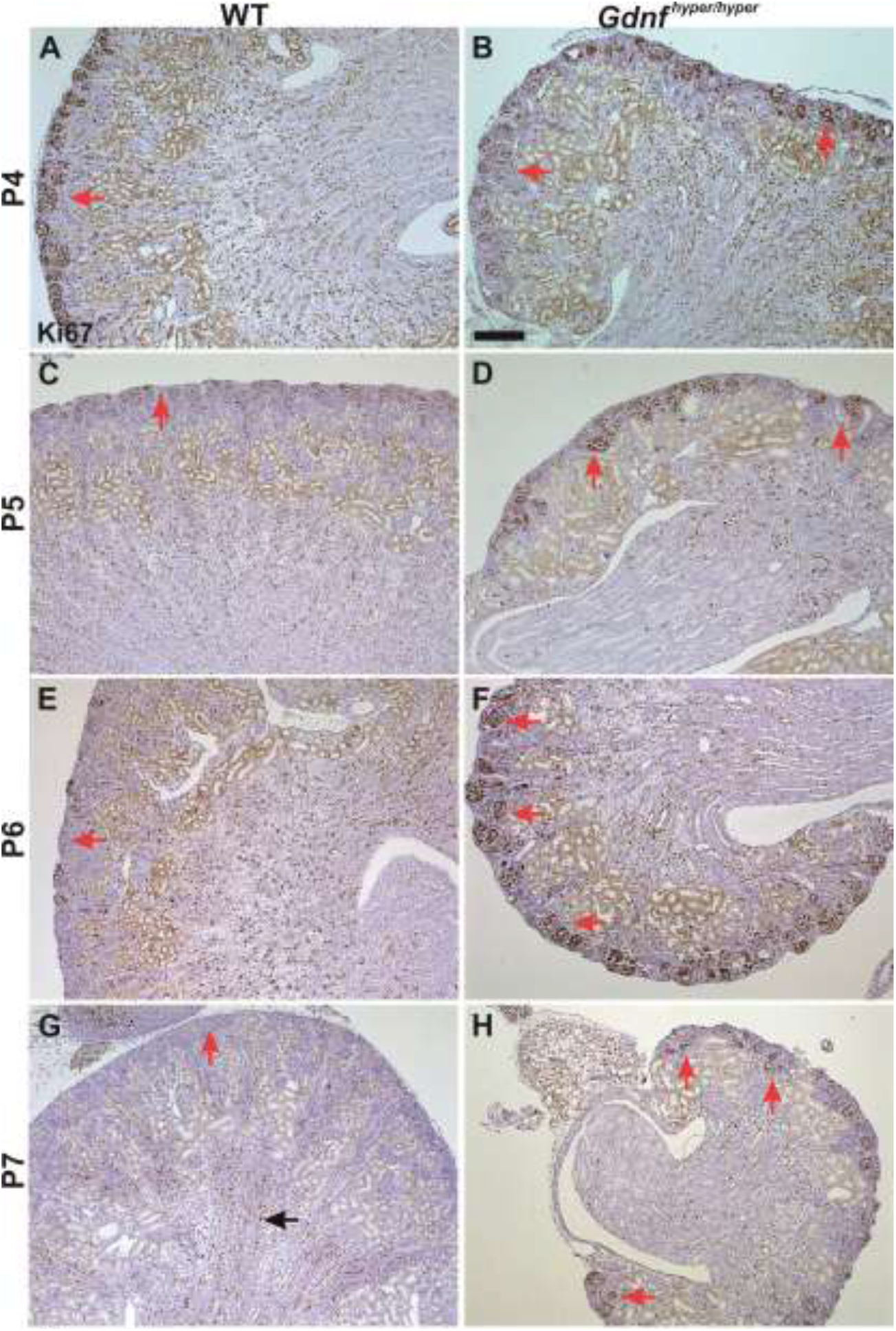
Sustained cortical proliferation in postnatal *Gdnf*^hyper/hyper^ kidneys. (A) Ki67 staining in P4 WT and (B) *Gdnf*^hyper/hyper^ kidneys shows similar pattern of proliferative cells in the cortical kidneys (red arrows) of both genotypes. (C) In P5 WT kidneys Ki67-positivity is significantly reduced in the cortex (red arrow) while (D) in *Gdnf*^hyper/hyper^ kidneys still exhibit focally very high proliferation (red arrows). (E) Proliferation in the WT cortical kidney is further diminished at P6 while maintained highly active in (F) *Gdnf*^hyper/hyper^ kidneys. (G) Ki67-positive proliferative cells localize to medullar renal tubules (black arrow) in P7 WT kidneys but (H) active proliferation is still detected in cortical differentiating nephron structures (red arrows) of *Gdnf*^hyper/hyper^ kidneys. *Scale bar:* 1mm.

Analysis of nephron precursors at P4-7 revealed an excessive quantity of LEF1, CYCLIN D1 and JAG1 positive nephron precursors in *Gdnf*^hyper/hyper^ kidneys until P7 while these were detected in control kidneys only up to P4 (Figure 6, Supplementary figure 4C-F). Such persistent postnatal nephrogenesis is able to maintain nephron density in *Gdnf*^hyper/hyper^ kidneys almost at the level of that in late embryonic kidneys (Supplementary figure 5). This demonstrates that a sustained postnatal NP pool in *Gdnf*^hyper/hyper^ kidneys is capable of inducing new nephrons after P3 and suggests that the prolonged nephrogenic program has functional significance for their survival.

**Figure 6.**
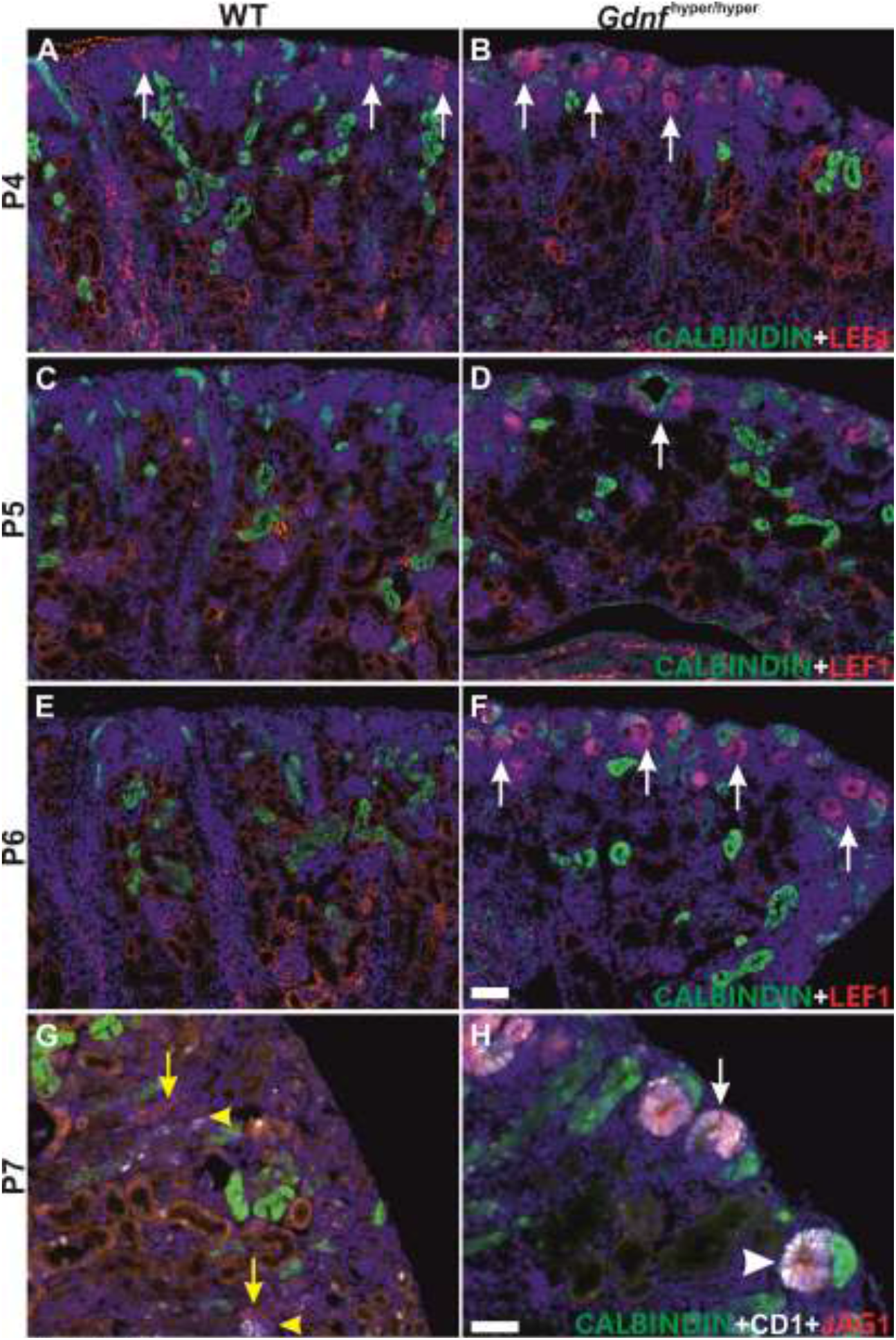
The distribution of nephron precursors in postnatal WT and *Gdnf*^hyper/hyper^ kidneys. (A) Nephron precursor marker LEF1 (red) in WT and (B) *Gdnf*^hyper/hyper^ kidneys at P4 demonstrates ongoing nephrogenesis in both genotypes (arrows). (C) P5 WT kidneys show very few LEF1 positive cells in cortex while ample nephrogenesis is detected in (D) *Gdnf*^hyper/hyper^ kidneys (arrows). (E) LEF1 is no longer present in the cortex of WT kidneys at P7 while plentiful nephron precursors are still detected in (F) *Gdnf*^hyper/hyper^ kidneys. (G) CYCLIN D1 (white) and JAG1 (red) localize to differentiated distal tubules of nephron (yellow arrow and arrowhead) in P7 WT kidney. (H) In P7 *Gdnf*^hyper/hyper^ kidneys CYCLIN D1 and JAG1 localize to comma-(arrowhead) and S-shaped (arrow) bodies of nephron precursors in the cortical differentiation zone thus showing sustained nephrogenesis at this late postnatal stage. *Scale bar*: 100μm.

*Gdnf* expressed by the nephron progenitor populations is lost in postnatal wild type kidneys by P2 (www.gudmap.org) while its mRNA and protein were still present in *Gdnf*^hyper/hyper^ kidneys as late as P7 (Figure 7A-D). Our results conclusively demonstrate a flexibility in the nephrogenic program, which can be prolonged with excess endogenous GDNF. This is particularly important as it shows that mouse kidneys bear an endogenous potential for inducing new nephrons longer than was previously thought (Brown et al., 2015, Hartman et al., 2007, Short et al., 2014, Togel et al., 2017). The results also suggest, unlike recent reports (Urbach et al., 2014, Yermalovich et al., 2019), that prolonged nephrogenesis can be uncoupled from increased cancer risk as *Gdnf*^wt/hyper^ mice with up to 4-fold increase in *Gdnf* expression have normal lifespan (n= 62) and lack tumors in kidneys and other organs (Supplementary figure 6) and *Gdnf*^hyper/hyper^ mice show no signs of neoplasia or tumorigenesis (Kumar et al., 2015, Turconi, Kopra et al., 2020).

**Figure 7.**
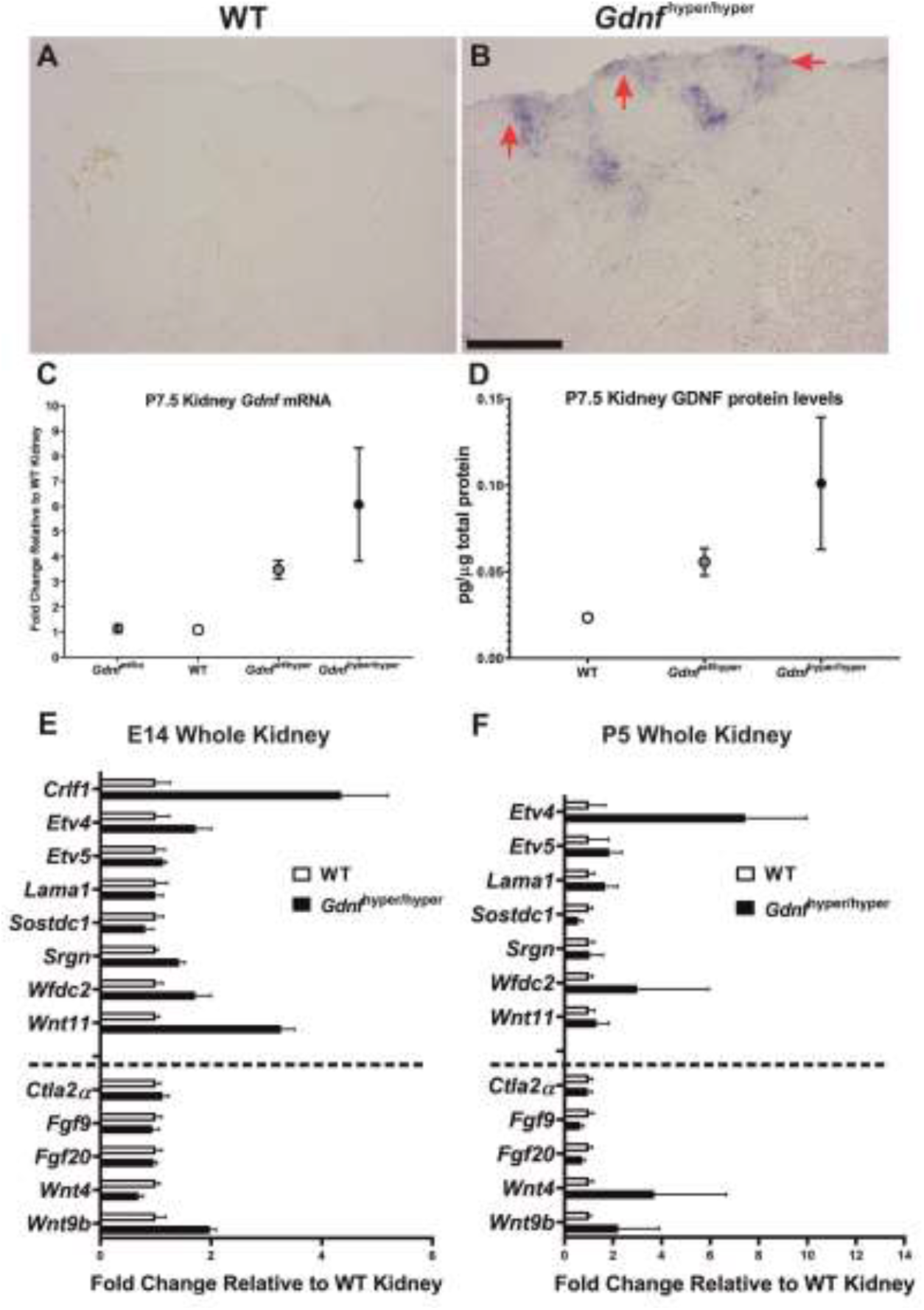
Expression analysis of the critical renal development regulators. (A) *In situ* hybridization assay of *Gdnf* at P4 is unable to detect any transcripts in wild type kidney. (B) *Gdnf* expression is maintained in P4 *Gdnf*^hyper/hyper^ kidney and localizes to cortical differentiation zone where nephron progenitors are maintained (red arrows) due to excess GDNF. (C) Quantitative RT-PCR (qRT-PCR) analysis of *Gdnf* transcripts in *Gdnf*^wt/ko^, WT, *Gdnf*^wt/hyper^ and *Gdnf*^hyper/hyper^ kidneys at P7. *Gdnf* expression in *Gdnf*^hyper/hyper^ kidneys is significantly higher than in WT (p=0.036) and *Gdnf*^wt/ko^ (p=0.038). (D) ELISA analysis of GDNF protein levels in P7 kidneys. (E) qRT-PCR based quantification of the relative expression levels of the selected genes at E14 and (F) at P5. The UB-derived secreted GDNF targets are listed above the dash line, while other known nephrogenesis regulators are presented below the line. *Gdnf*^hyper/hyper^ kidneys show significant upregulation of *Crlf1* (p=0.019), *Srgn* (p=0.021), *Wnt11* (p=0.001) and *Wnt9b* (p=0.01) at E14, while at P5, significantly upregulated genes include *Etv4* (p=0.000), *Etv5* (p=0.03), *Lama1* (p=0.014) and *Wnt4* (p=0.026). In contrast, the transcription of Sostdc1 (p=0.000), Fgf9 (p=0.001) and Fgf20 (p=0.001) is significantly downregulated in *Gdnf*^hyper/hyper^ kidneys at P5. *Scale bar*: 100μm.

### Persistent postnatal nephrogenesis depends on GDNF induced signaling changes

The conservative receptor complex for GDNF is composed of the RET receptor tyrosine kinase and the co-receptor GFRα1 (Costantini, 2010) co-expressed only in the epithelial UB tips (Golden, DeMaro et al., 1999, Pachnis et al., 1993). We reasoned that the effects of increased endogenous GDNF on nephrogenesis must be mediated through GDNF-regulated (Lu et al., 2009, Ola et al., 2011), UB-derived secreted factor(s) (http://www.signalpeptide.de/index.php) (Schmidt-Ott et al., 2005). Analysis of known GDNF targets identified eight potential candidates matching to these criteria (Table 1).

**Table 1.** Secreted GDNF target genes with potential to influence nephrogenic program. The table indicates in which screen the downstream target gene was identified (Lu et al., 2009, Ola et al., 2011) and experimental evidence for responsiveness to GDNF. Asterisk marks those genes that have been recognized as secreted factors that are potentially involved in nephrogenesis regulation (Schmidt-Ott et al., 2005).

| ***Lu et al*** |  |  |
| --- | --- | --- |
| <u>Gene</u> | <u>GDNF response</u> |  |
| Crlf1* | + | Cytokine receptor like factor 1 |
| Sostdc1* | + | Sclerostin domain-containing protein 1 |
| Wnt4 | - | Wingless-type MMTV integration site family, member 4 |
| Wnt11 | + | Wingless-type MMTV integration site family, member 11 |
| ***Ola et al*** |  |  |
| Crlf1 | + | Cytokine receptor like factor 1 |
| Srgn | + | Serglycin |
| Wfdc2* | + | WAP four-disulfide core domain 2 |
| Ctla2a | + | cytotoxic T lymphocyte-associated protein 2 alpha |
| Lama1* | + | Laminin, subunit alpha 1 |

The expression analysis of UB-derived secreted GDNF-targets revealed significant upregulation in *Crlf1*, *Srgn* and *Wnt11* at mid-gestation while they all were normalized in postnatal *Gdnf*^hyper/hyper^ kidneys (Figure 7E-F). This indicates not only increased GDNF-induced signaling as demonstrated by increased target transcription factor *Etv4* and *-5* expression(Kuure, Chi et al., 2010, Lu et al., 2009) (Figure 7E-F), but also demonstrates augmented reciprocal communication from mutant UB to the adjacent NP pool. Also, upregulation of *Wnt9b*, known to regulate the balance between NP cell maintenance versus differentiation (Carroll et al., 2005, Karner et al., 2011, Kiefer et al., 2012), and downregulation of the nephrogenesis inducer *Wnt4*, were detected in *Gdnf*^hyper/hyper^ embryonic kidneys while *Wnt4* expression was reversed in postnatal kidneys (Figure 7E-F).

We next functionally tested if the most significant gene expression changes in *Gdnf*^hyper/hyper^ kidneys, namely *Crfl1* and *Wnt11*, affect the nephrogenic program in the mouse. Cytokine receptor-like factor 1 (*Crlf1*, also known as CLF-1) is a UB tip specific gene, which in a complex with its physiological ligand cardiotrophin-like cytokine (CLC) induces isolated kidney mesenchyme for differentiation in rat (Schmidt-Ott et al., 2005). The characterization of *Crfl1* knockout kidneys revealed kidney size and nephrogenesis comparable to the wild type kidneys (Supplementary figure 7A-D, n=6), suggesting that despite its capacity to trigger differentiation in NP pools, the *in vivo* function of CRLF1 is redundant either with other CRLF family members or with other molecules regulating the nephrogenic program (O’Brien, 2018, Schmidt-Ott et al., 2005).

WNT11 coordinates ureteric bud branching and was recently shown to regulate NP maintenance and ultimately nephron endowment (Majumdar, Vainio et al., 2003, Nagy, Xu et al., 2016, O’Brien et al., 2018). To investigate whether significantly increased *Wnt11* expression in *Gdnf*^hyper/hyper^ kidneys contributes to the altered nephrogenic program in *Gdnf*^hyper/hyper^ mice they were crossed to the *Wnt11* knockout background (Majumdar et al., 2003). This failed to generate double homozygote *Gdnf*^hyper/hyper^;*Wnt11*^−/−^ offspring (n=95) from the compound heterozygote (*Gdnf*^wt//hyper^;*Wnt11*^+/−^) breeding. A statistically significant deviation from Mendelian ratios was confirmed for divergent gene distribution by χ^2^ test (p=0.005, Table 2). These results indicate embryonic lethality for *Gdnf^h^*^yper/hyper^;*Wnt11*^−/−^ mutants, and prevent the analysis of double mutant kidneys. Therefore we focused our analysis on *Gdnf^h^*^yper/hyper^;*Wnt11*^+/−^ kidneys.

**Table 2.** Summary of offspring genotypes from *Gdnf^wt//hyper^;Wnt11*^+/−^ intercrosses. The offspring genotyping analysis was performed between E14.5 and newborn stages and it failed to identify any individual with *Gdnf^h^*^yper/hyper^;*Wnt11*^−/−^ genotype. Statistical analysis with χ^2^ test reveals that gene distribution significantly deviates from Mendelian ratio (P=0.005).

|  | *Wnt11*<sup>wt/wt</sup> | *Wnt11*<sup>wt/-</sup> | *Wnt11*<sup>-/-</sup> |
| --- | --- | --- | --- |
| *Gdnf*<sup>wt/wt</sup> | 6 | 13 | 2 |
| *Gdnf*<sup>wt/hyper</sup> | 26 | 22 | 2 |
| *Gdnf*<sup>hyper/hyper</sup> | 9 | 15 | 0 |

Removal of one functional *Wnt11* allele in *Gdnf*^hyper/hyper^ background (*Gdnf*^hyper/hyper^*;Wnt1^+/−^*) did not improve postnatal kidney size or morphology from that in *Gdnf*^hyper/hyper^ (Supplementary figure 8A-E). Importantly, genetic *Wnt11* reduction in *Gdnf^h^*^yper/hyper^ embryos resulted in expansion of SIX2-positive NP cells, which was further supported by PAX2 staining, thus demonstrating amelioration of the nephrogenic program in *Gdnf*^hyper/hyper^*;Wnt1*^+/−^ animals (Figure 8A-H, compare also to Figure 1F).

**Figure 8.**
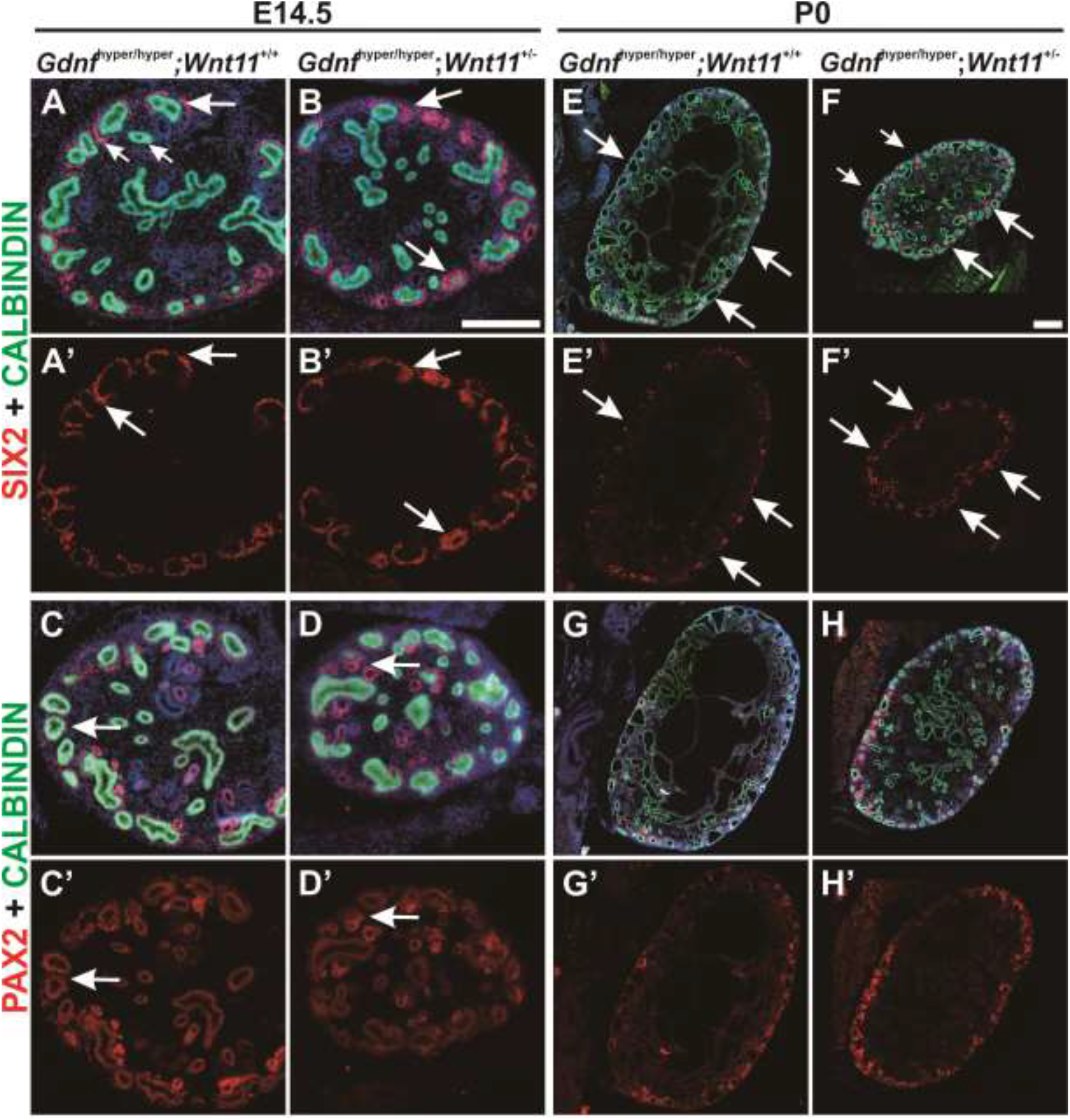
Genetic reduction of *Wnt11* level partially rescues nephron progenitor loss in *Gdnf*^hyper/hyper^ kidneys. (A) Localization of SIX2-positive nephron progenitors (red) in E14.5 *Gdnf*^hyper/hyper^ and (B) *Gdnf*^hyper/hyper^; *Wnt11*^+/−^ kidneys shows increment in general abundance of nephron progenitors per niche in the absence of one *Wnt11* allele. (C) PAX2-positive cells in *Gdnf*^hyper/hyper^ and (D) *Gdnf*^hyper/hyper^; *Wnt11*^+/−^ kidneys show similar patterns and amount at E14.5. (E-H) At P0 kidneys both SIX2 and PAX2 staining demonstrate improvement in the absence of one *Wnt11* allele. Distribution of (A’, B’, E’, F’) SIX2-positive nephron progenitors and (C’, D’, G’, H’) PAX2-positive cells are shown without CALBINDIN (green) signal, which depicts the UB epithelium in all kidneys. *Scale bar*: 200μm.

## DISCUSSION

Each person’s nephron count is defined during his/her fetal life and can vary greatly between different individuals. Epidemiological studies indicate that those born with a higher nephron count are better protected against kidney disease and have a lower risk for hypertension and cardiovascular diseases later in life (Starr & Hingorani, 2018). Neonephrogenesis, the formation of new nephrons after the cessation of the fetal nephrogenic program observed in some vertebrates, is not an inherent feature of mammalian kidneys (Camarata, Howard et al., 2016). When diseased or injured, mammalian nephrons permanently lose their function as the adult kidney shows very limited regeneration, and instead responds to insult by compensatory growth through glomeruli and tubular hypertrophy (Chawla & Kimmel, 2012, Romagnani et al., 2013, Westland, Schreuder et al., 2014). Thus, identification of the mechanisms controlling nephron endowment is essential for the development of means to safely maximize nephron endowment and thereby renal health.

Here we show that the neurotropic factor GDNF, known to initiate renal development and regulate ureteric bud morphogenesis (Costantini, 2010, Kurtzeborn et al., 2018, Li et al., 2019), is capable of prolonging the nephrogenic program beyond its normal cessation. The complete mechanism through which GDNF does this remains to be detailed, but our results demonstrate that excess GDNF has a major effect on nephron progenitor (NP) self-renewal and maintenance. Initially in embryonic kidneys, high GDNF levels cause a rapid decline in NP numbers per niche, which is typical for many mouse models with primary deficiency in ureteric bud (UB) that is needed for normal NP niche composition (Brown et al., 2015, Carroll & Das, 2013, Chen, Yao et al., 2015b, Costantini & Kopan, 2010, Liu, Chen et al., 2018, O’Brien & McMahon, 2014). The premature NP decline in these previously published models is caused by their untimely differentiation, which is not the cause in *Gdnf*^hyper/hyper^ kidneys. Instead, excess GDNF is capable of maintaining an active cell cycle in the postnatal kidney cortex significantly longer than in wild type kidneys. Importantly, proliferation occurs in NP cells, which are maintained longer than in wild type kidneys and keep producing new nephrons with a higher rate and beyond the previously defined cessation point (Hartman et al., 2007, Rumballe et al., 2011, Short et al., 2014). These results indicate that GDNF can prolong the lifespan of NPs with favorable consequences on final nephron count.

Our data demonstrate that the decline in embryonic NPs of *Gdnf*^hyper/hyper^ kidneys is due to decreased cell proliferation rather than increased premature differentiation. This cellular mechanism is supported by molecular changes in known GDNF targets (*Crlf1*, *Wnt11* and *Srgn*) (Lu et al., 2009, Ola et al., 2011) and in important regulators of NP cell maintenance (*Wnt9b* and *Wnt11*) (Karner et al., 2011, Kiefer et al., 2012, O’Brien et al., 2018). Together with previous publications specially addressing longevity and resilience of NP cells by ablating the NP population with diphtheria toxin A or disturbing MAPK/ERK activation, which both resulted in fast decline in NP cells without positive effect on their lifespan (Cebrian et al., 2014, Ihermann-Hella, Hirashima et al., 2018), this suggests that NP cell proliferation inherently plays a major role in defining their total lifetime. In agreement, NP cells in late embryonic kidneys show significantly reduced proliferation rates and accelerated differentiation, while the transplantation of old NP cells into earlier stage kidney prolongs their lifespan (Chen et al., 2015a).

We propose that a mechanism through which NP cells sense their developmental age and ultimately lifespan is related to GDNF, and possibly other signaling activity levels (see below). In a normal kidney, GDNF expression is high at the initiation of kidney morphogenesis and during the active growth phase, while a clear decline in GDNF levels occurs in late embryonic kidney resulting in loss of expression by early postnatal stage (Fig. 7). GDNF levels in early *Gdnf*^hyper/hyper^ are abnormally high, which appears inhibitory for NP cell proliferation, likely due to the cellular and molecular changes within mutant UB. Similar to wild type kidneys, GDNF levels decrease in postnatal *Gdnf*^hyper/hyper^ kidneys as the endogenous control of gene expression is maintained intact(Kumar et al., 2015), and this diminishes GDNF expression to the level that is permissive for NP cell self-renewal and differentiation, which are detected as a sustained nephrogenic program in postnatal kidneys. Such a newly recognized, growth factor-dependent plasticity in renal differentiation bears remarkable hope for its utilization in regenerative purposes.

The first clues for the possibility to lengthen the nephron differentiation period came from experiments inhibiting BMP/SMAD1/5 signaling, which led to slightly bigger kidneys with more nephrons than in vehicle treated kidneys (Brown et al., 2015). Decreased dosage of *Hamartin* (*Tsc1*), encoding an mTOR inhibitor, maintains NP cells for an additional day, without reported effects on embryonic NP cell numbers (Volovelsky et al., 2018). Finally, prolonged nephrogenesis was observed also in mice with overexpression of *Lin28*, *-28b* or inactivated *Let-7* (Urbach et al., 2014, Yermalovich et al., 2019). Despite their great potential, these models show either direct tumorigenesis or are associated with increased activation of Igf2/H19 locus, which associates with pediatric kidney cancer known as Wilms’ tumor (Chen, Stroup et al., 2018, Ogawa, Eccles et al., 1993). As a note, no changes in pSMAD1/5 localization or levels were detected in *Gdnf*^hyper/hyper^ kidneys, which are tumor-free also in aged heterozygotes animals.

The present study focuses on the characterization of the GDNF’s function in nephrogenesis, which has been previously examined by diminishing endogenous *Gdnf* dosage that reduces nephron endowment (Cullen-McEwen, Drago et al., 2001, Cullen-McEwen, Kett et al., 2003). The canonical receptor complex for GDNF is formed by the RET tyrosine kinase and its co-receptor GFRa1, which are exclusively co-expressed only in the UB epithelium (Costantini, 2012, Golden et al., 1999, Pachnis et al., 1993). We identified UB-derived, secreted CRLF1 and WNT11 as potential GDNF signaling targets that mediate its effects on NP regulation. Our data suggests that the function of CRLF1 is redundant with other cytokines as its inactivation does not affect renal differentiation. WNT11 has been demonstrated crucial for the maintenance of normal nephrogenesis (Majumdar et al., 2003, Nagy et al., 2016, O’Brien et al., 2018), and its expression is positively regulated by GDNF (Costantini & Shakya, 2006, Majumdar et al., 2003, Sainio, Suvanto et al., 1997). Importantly, lowering *Wnt11* expression in the *Gdnf*^hyper/hyper^ background clearly improved NP maintenance and differentiation, but failed to fully rescue the renal hypoplasia caused by the excess GDNF. This may at least partially derive from the failure to inactivate both *Wnt11* alleles in the *Gdnf*^hyper/hyper^ background, which likely is due to the previously reported embryonic lethality of *Wnt11*^−/−^ in the SV129 background (Nagy et al., 2016). Thus, one remaining *Wnt11* allele may even augment the GDNF cascade as suggested earlier (Majumdar et al., 2003). It is probable that WNT9b, expressed by the UB and known to regulate many aspects of nephrogenesis (Carroll et al., 2005, Karner et al., 2011, Kiefer et al., 2012), also participates in mediating GDNF’s effects on nephrogenesis as its expression was greatly affected by the high GDNF dosage.

The current study cannot rule out the possibility that some of the GDNF’s effects on NPs are due to non-canonical GDNF signaling as its potential alternative signaling receptors are expressed by the NPs (Brodbeck, Besenbeck et al., 2004, Enomoto, Hughes et al., 2004, Keefe Davis, Hoshi et al., 2013, Paratcha, Ledda et al., 2003). However, both deletion of NCAM or expression of *Gfrα1* in UB only (by replacing it under the control of *Ret* promoter) maintain a normal renal phenotype (Cremer, Lange et al., 1994, Heuckeroth, Enomoto et al., 1999, Keefe Davis et al., 2013, Rossi, Luukko et al., 1999, Sariola & Saarma, 2003). This suggests that at least on their own, neither NCAM nor GFRa1 are critically required for the nephrogenic program.

Our results show that prolonged preservation of NPs in postnatal kidneys sustains postnatal nephrogenesis in the *Gdnf*^hyper/hyper^ kidneys. First, the re-synchronization of the nephrogenesis pace to that in wild type kidneys is followed by a remarkable extension of the nephrogenic program. This, unlike in the *Gdnf* inactivation model, allows a big enough nephron endowment for *Gdnf*^hyper/hyper^ mice to survive the postnatal lethality. Thus, GDNF, which has been and currently is in clinical trials for Parkinson’s disease, may bear potential to serve as a prospective mean to safely improve nephron endowment in prematurely born infants who suffer from reduced nephron counts(Starr & Hingorani, 2018).

## METHODS

### Animals

Animal care and research protocols were performed in accordance with the Code of Ethical Conduct for Animal Experimentation as well as European Union directives (Directive 2010/EU/63), and were approved National Animal Experiment Board of Finland.

Mice were housed in individually ventilated cages with food and water *ad libitum*. Optimal humidification and heating were provided constantly at the certified Laboratory Animal Centre of University of Helsinki.

The generation and genotyping of the *Gdnf^hyper^* (Kumar et al., 2015) and *Fgf9;20* (Barak et al., 2012) mouse models has been described previously. *Gdnf^hyper^;Wnt11*^−^ pups are from intercrosses of *C57BL6* Wnt11 ^−^(Nagy et al., 2016) and *Gdnf^hyper^* lines, and were genotyped as previously described. All the lines used in the studies were maintained on a 129Ola/ICR/C57bl6 triple mixed background.

The generation of *Crlf* knockout mice is described previously (Alexander, Rakar et al., 1999). PCR genotyping was performed using a Neo specific primer set (5’AGAGGCTATTCGGCTATGACTG3’ and 5’CCTGATCGACAAGACCGGCTTC3’) together with gene specific primer sets for *Crlf1* (5’GCCTAATAGGTGCTGGGTGA 3’ and 5’ GACCCTATCTGCGTTTTCCA 3’).

### Tissue collection

Embryos were staged according to the criteria of Theiler (Theiler, 1989) as described previously(Kuure, Sainio et al., 2005) and collected after cervical dislocation of pregnant females at deep anesthesia with CO_2_. Late embryonic and postnatal pups were sacrificed via decapitation. Tissue was dissected in *Dulbecco*’*s* medium supplemented with 0.2% bovine serum albumin followed by fixation with 4% PFA or organ culture.

### Organ culture

Kidneys isolated from E11.5 and E12.5 mouse embryos were placed at the air-liquid interface supported by Transwell^®^ polyester membrane system (Costar) and cultured at 37 °C in a humidified 5% CO_2_ atmosphere with kidney culture medium (F12:DMEM (1:1) + Glutamax (Gibco) supplemented with 10% fetal bovine serum and penicillin-streptomycin)(Ihermann-Hella & Kuure, 2019). For the *in vitro* culture with exogenous GDNF (100 ng/mL, ProSpec Ltd), each urogenital block containing metanephros, the definitive kidney rudiment, was halved along the *linea mediana ventralis*. One half of each sample was cultured with exogenous GDNF while the other half was served as the control.

### Histology and immunostaining

For studies investigating the marker of interest on paraffin section, kidneys were dissected from the embryos or pups of the indicated stages, followed by tissue processing and sectioning as referred to in previous publication (Li et al., 2019). Hematoxylin-eosin (H&E) and immunohistochemistry performed on paraffin sections were completed as previously described(Li et al., 2019). In brief, the dissected tissue was fixed overnight with 4% PFA (pH 7.4), followed by dehydration and paraffinization with an automatic tissue processor (Leica ASP 200). Sections with the thickness of 5 μm were prepared and dewaxed in Xylene followed by rehydration before being stained with Harris hematoxylin and Eosin or undergoing heat-induced antigen retrieval in antigen retrieving buffer (10 mM Citrate; pH 6.0). For immunostaining, 3% BSA was used for blocking (1 hour at room temperature) followed by 30 min 0.5% hydrogen peroxide treatment for chromogenic staining. Sections were then incubated with primary antibodies overnight at + 4 ℃ followed by species-specific secondary antibody incubation for 2h at room temperature. Chromogenic detection was implemented with EnVision Detection System-HRP (DAB) kit (Dako).

Tissues subjected to whole-mount immunofluorescence staining were fixed and permeabilized with 100% ice-cold methanol for 10 minutes before overnight incubation with primary antibodies. Incubation with the corresponding species-specific secondary antibodies was implemented on the second day following the vigorous wash with PBST (0.1% Tween 20 in PBS).

The information about the primary and secondary antibodies used in the present study is listed in Supplementary Table 1.

Any conclusion based on immunostaining and histology were obtained via the observation of multiple kidneys (n≥3/each genotype) in order to evade bias.

### *In situ* hybridization

In situ hybridization was conducted as previously described (Ihermann-Hella, Lume et al., 2014). In brief, anti-sense RNA probe against *Gdnf* exons was transcribed and hybridized on thick sections derived from P4 kidneys embedded in 4% low melting agarose (NuSieve GTG, Lonza). BM purple was used for colorimetric reaction.

### ELISA

GDNF protein levels in P7.5 kidneys were measured via ELISA as reported previously (Kumar et al., 2015). GDNF Emax^®^ Immunoassay (Promega) was utilized with acid treatment when carrying out the ELISA.

In detail, for each genotype, 3 kidneys were lysed immediately after being dissected. After measuring the protein concentration with DC protein Assay (Bio-Rad), 25 μg of total protein was loaded into the well on ELISA plate. Each sample was analysed in duplicate.

### Imaging

Immunostaining and *in situ* hybridization on sections were imaged either with Zeiss AxioImager equipped with the HXP 120 V fluorescence light source, Hamamatsu Orca Flash 4.0 LT B&W camera and Zeiss AxioCam 105 color camera, or with Leica DM5000B equipped with Leica EL 6000 metal halide light source and Hamamatsu Orca-Flash4.0 V2 sCMOS camera. Whole mount immunofluorescence staining was imaged with Zeiss AxioImager with the Zeiss ApoTome application.

### Reverse Transcription and Quantitative RT-PCR (qRT-PCR)

qRT-PCR experiments were conducted as reported previously (Kumar et al., 2015). More detailed, 150 to 500 ng of total RNA from each sample was treated with RNase-free DNase I (Thermo Fisher Scientific). The reverse transcription reaction was performed with random hexamer primers and RevertAid Reverse Transcritase (Thermo Fisher Scientific) immediately after inactivation of DNase I with 5 mM EDTA at 65 °C. Complementary DNA (cDNA) was diluted 10 times and stored at - 20 °C until the conduction of qRT-PCR.

For qRT-PCR, LightCycler 480 SYBR Green I Master (Roche) or Bio-Rad C1000 Touch Thermal Cycler upgraded to CFX384 System (Bio-Rad), supplied with SYBR Green I Master (Roche) and 250 pmol primers was loaded into the well of 384-well plates with a total volume of 10 μL. Negative control was always included in every reaction via setting conditions with minus-reverse transcription or water. A combination of *Pgk1*, *Hprt* and *Gapdh* were used as reference genes in all the qRT-PCR experiments except the one with P7.5 kidneys, in which mouse *Actβ* was selected as reference gene.

Data obtained from quantitative RT-PCR were analyzed as described previously (Kumar et al., 2015). Normalization was performed according to the geometric mean of the reference genes. All samples were analysed in a duplicate manner. Results for a biological repeat were discarded when the Cq value for one or more of the replicates was 40 or 0, or when the Cq difference between replicates was >1.

Sequences of the primer used can be found in Supplementary Table 2.

### Quantitative Analyses

Quantification of the density of glomeruli was calculated with the sequential sections of P0 and P7 kidneys. The sections were selected every 125 μm to avoid iteration count of glomeruli. The number of glomeruli and the area of each kidney on the selected sections were counted manually with Fiji ImageJ software (V1.51) on each section selected. Only glomeruli with intact shape and can be distinguished from were recorded. The volume of renal mass is calculated with the equation 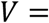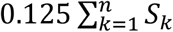, the area of each kidney section. In order to compensate the number of omitted nephrons, the quantity of total glomeruli within the investigated renal mass is calculated with the equation 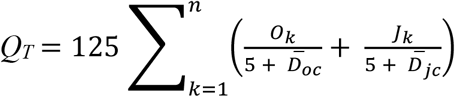. O_k_, the number of glomeruli in outer cortical region of each kidney section; J_k_, the number of glomeruli in juxtamedullary cortical region of each kidney section; 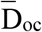, the average diameter of the biggest glomeruli in the outer cortical region shown on each section from one kidney (μm); 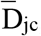, the average diameter of the biggest glomeruli in the juxtamedullary cortical region shown on each section from one kidney (μm). Finally, the density of glomeruli in each kidney is calculated with the equation 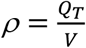.

For quantification of nephron progenitors, the images captured with the application of Zeiss ApoTome were processed with Imaris Software (Bitplane) using spot tracking. All images were processed with exactly identical protocols for analysis.

### Statistical Analysis

All values are presented as mean ± standard error of the mean (SEM) except the main effects analysis of *Gdnf* and *Wnt11* on kidney size, which are shown as mean ± standard deviation (SD). Statistical significance was set at *p* < 0.05 for all the analysis.

Independent-samples two-tailed *t* test was utilized for pairwise comparisons with SPSS Statistics software (IBM; Version 25) in the study unless otherwise noted. The data obtained via *in vitro* tissue culture with exogenous GDNF was analyzed with paired-sample *t*-test using SPSS Statistics software (IBM; Version 25). The impacts of genetically modified *Gdnf* and *Wnt11* expression on kidney size were analyzed via Two-way ANOVA followed by main effects test with SPSS Statistics software (IBM; Version 25). The statistical significance for the deviation of gene distribution from Mendelian ratio was performed utilizing χ^2^ test also with SPSS Statistics software (IBM; Version 25). The data obtained from ELISA and qRT-PCR of P7.5 kidneys were analyzed with One-way ANOVA followed by Bonferroni *post-hoc* test using SPSS Statistics software (IBM; Version 25).

## ACKNOWLEDGEMENTS

We thank MSc Kristen Kurtzeborn for fruitful discussions and critical reading of the manuscript, and the personnel of Biomedicum Imaging and Light Microscopy Units at of Helsinki Institute of Life Sciences (HiLIFE) for their valuable help in image analysis and quantification related techniques. The mice were housed at Laboratory Animal Centre of HiLIFE, University of Helsinki. S.K. was supported by grants from the Academy of Finland (309997), Jane and Aatos Erkko, and Maud Kuistila Foundations, and H.L. by the Doctoral program in Biomedicine, University of Helsinki. J.O.A. was supported by the Academy of Finland (grant no. 297727), Sigrid Juselius Foundation, Faculty of Medicine at the University of Helsinki, Helsinki Institute of Life Science (HiLIFE) Fellow grant, European Research Council (ERC) (grant no. 724922).

## AUTHOR CONTRIBUTIONS

S.K. together with H.L. and J.O.A. contributed to the design of the project. Experimental work was carried out by H.L., J.K, Y.G., E.S., B.L, K.M, S.O. and A.R.M.-R. Analysis of results, their interpretation and presentation was carried out by H.L. and S.K. with additional contribution from J.O.A., S.-H.H. and F.C.

## SUPPLEMENTARY FIGURE LEGENDS

**Supplementary figure 1 - Decreased NP proliferation in *Gdnf*^hyper/hyper^ kidneys is caused by high level of GDNF in early embryonic kidneys.** (A) Three-dimensional reconstructed apotome z-stacked images of representative E11.5 WT and (B) *Gdnf*^hyper/hyper^ kidneys stained with E-CADHERIN (yellow), CALBINDIN (yellow) and SIX2 (green). (C) Comparison of SIX2-positive NP amount in WT and *Gdnf*^hyper/hyper^ kidneys shows increased initial NP pool in *Gdnf*^hyper/hyper^ kidneys at E11.5. Data are presented as mean quantity of SIX2-positive NPs ± SEM compared with those of WT littermates, which are set to 100% and reflect the results obtained from four independent litters (WT: 100 ± 12.15 %, n=7; *Gdnf*^hyper/hyper^: 163.59 ± 6.42 %, n=9; P=0.000). (D) Similar analysis at E12.5 indicates that the quantity of NP in *Gdnf*^hyper/hyper^ kidneys is normalized to that in WT kidneys. Data are presented as mean quantity of SIX2-positive NPs ± SEM compared with those of WT littermates, which are set to 100% and reflect the results obtained from three independent litters (WT: 100 ± 15.61 %, n=6; *Gdnf*^hyper/hyper^: 124.74 ± 33.89 %, n=5; P=0.533). (E) Proportion of proliferating SIX2-positive NPs in *Gdnf*^hyper/hyper^ kidneys at E12.5 shows a significant decrease when compared to wild type kidney. Data are presented as mean proportion of proliferating SIX2-positive NPs ± SEM compared with those of WT littermates which are set to 100% and reflect the results obtained from three independent litters. (WT: 100 ± 9.04 %, n=5; *Gdnf*^hyper/hyper^: 56.94 ± 12.67 %, n=5; P=0.024). (F) Proportion of proliferating SIX2-positive NPs in E11.5 kidneys cultured for 24 hours with or without (control) 100 ng/mL GDNF supplementation. (Control: 6.25 ± 0.34 %, n=5; GDNF: 4.64 ± 0.33 %, n=5; P=0.012). * denotes P < 0.05, *** denotes P < 0.001; unpaired 2-tailed *t*-test. *Scale bar*: 30μm.

**Supplementary figure 2 - PAX2 staining in embryonic wild type and *Gdnf*^hyper/hyper^ kidneys.** (A) PAX2 (red) and E-CADHERIN (green) localization in wild type (WT) and (B) *Gdnf*^hyper/hyper^ kidneys at E11.5. (C) PAX2 (red) and CALBINDIN (green) staining in WT and (D) *Gdnf*^hyper/hyper^ kidneys at E14.5 shows decrease in nephron progenitors (arrows) and enlargement of ureteric bud epithelium (green). Asterisks mark the differentiating nephron precursors, which are greatly reduced in *Gdnf*^hyper/hyper^ kidneys. *Scale bar*: 100μm.

**Supplementary figure 3 - Nephron progenitors are not maintained in postnatal kidneys upon *Fgf9* and *Fgf20* deletion.** Representative images showing SIX2 (red) and E-CADHERIN (green) staining in (A) *Fgf20*^+/−^, (B) *Fgf20*^−/−^ and (C) *Fgf9*^+/−^;*Fgf20*^/-^ kidneys at P6. (D) PAX2 (red) and E-CADHERIN (green) staining in *Fgf20*^+/−^, (E) *Fgf20*^−/−^ and (F) *Fgf9*^+/−^;*Fgf20*^−/−^ kidneys at P6 all indicate similar result. Cap mesenchyme that would be positive for SIX2 or PAX2, indicating the presence of nephron progenitors and differentiating precursors, cannot be detected in any of the FGF-manipulated kidneys. *Scale bar*: 50μm.

**Supplementary figure 4 - Sustained cortical proliferation in postnatal nephrogenesis in *Gdnf*^hyper/hyper^ kidneys.** (A) Ki67 staining visualizes similar pattern of proliferative cells in P12 WT and (B) *Gdnf*^hyper/hyper^ kidneys. (C) Co-staining of E-CADHERIN (green) and Ki67 (red) in P4 WT and (D) *Gdnf*^hyper/hyper^ kidneys. White arrows point to nephron precursors, which are double positive for Ki67 and E-CADHERIN, red arrows indicate newly induced renal vesicles, which not yet have acquired the expression of E-CADHERIN. Yellow arrows point to distal tubules of nephrons positive for Ki67 and abnormally positioned to renal cortex. Arrowheads indicate ureteric buds, which also show proliferative activity. (E) P4 WT kidney stained with JAG1 (red), CALBINDIN (green) and Ki67 (white) shows proliferation in differentiating nephron precursors (arrows). (F) Corresponding triple staining in *Gdnf*^hyper/hyper^ kidneys demonstrates slightly more on-going nephrogenesis (arrows) but similar proliferation pattern. *Scale bar*: 100μm.

**Supplementary figure 5 - Glomerular density assessment.** Representative images showing the distribution of glomeruli in P7 WT kidney (A) and *Gdnf*^hyper/hyper^ kidneys (B). Chromogenic antibody staining was performed for PODOCALYXIN and counterstained with hematoxylin. (C) Quantification of nephron density in *Gdnf*^hyper/hyper^ kidneys at E18.5 and P7. Data are presented as the mean quantity of nephron per cubic millimeter ± SEM. (E18.5: 131 ± 10.00, n=4; P7: 119.99 ± 13.67, n=4; P=0.507). *Scale bar*: 1mm.

**Supplementary figure 6 – Elevated endogenous *Gdnf* expression level does not promote tumorigenesis.** (A) Representative image of adult *Gdnf^wt/hyper^* mouse (five months) with thoracic cavity and peritoneal cavity exposed. No tumours were observed in any organs based on the anatomical examination (n=12). (B) Representative image of HE stained kidney section from adult *Gdnf^wt/hyper^* mouse shows no signs of neoplasia. Abbreviations: B; bladder, H; heart, L; liver, St; stomach, Sv; seminal vesicle. Scale bar: 1 mm.

**Supplementary figure 7 - Function of CRLF1 is dispensable for nephrogenesis**. (A) Analysis of kidney morphology with hematoxylin and eosin staining at P0 WT and (B) *Crlf1*^−/−^ kidneys (n=15) revealed normal kidney size and organization. (C) PODOCALYXIN immunofluorescence visualizes developing glomeruli in E11.5 kidneys cultured for 4 days from the WT and (D) *Crlf*1^−/−^ embryos. The average number of nephrons was 34 +/− 10 in CLF-1 −/− vs. 29 +/− 8 in wild type kidneys (p=0.34). *Scale bar*: 100 μm.

**Supplementary figure 8 - GDNF and WNT11 affect kidney size independently from each other.** (A) Main effect analysis of genetically increased GDNF on kidney size in *Gdnf*^hyper^;*Wnt11* compound background. Data are presented as estimated marginal means of area obtained via measuring the coronal plane of each kidney ± SD compared to those of *Gdnf*^wt/wt^;*Wnt11*^+/+^ controls and reflect the results obtained from five independent litters. The (mean) size of *Gdnf*^wt/wt^;*Wnt11*^+/+^ kidney(s) in each litter was set to 100% in order to normalize the difference among litters. (B) Main effect analysis of genetically decreased WNT11 on kidney size in *Gdnf*^hyper^;*Wnt11* compound background. Similar analysis was performed as in A. (C) Presentation of size analysis as whole. Data are presented as means of area obtained by measuring the coronal plane of each kidney ± SEM in comparison to *Gdnf*^wt/wt^;*Wnt11*^+/+^ controls, which are set to 100% and reflect the results obtained from five independent litters. (*Gdnf*^wt/wt^;*Wnt11*^+/+^: 100 ± 4.09%, n=6; *Gdnf*^wt/wt^;*Wnt11*^+/−^: 92.61 ± 3.53%, n=8; *Gdnf*^wt/wt^;*Wnt11*^−/−^: 58.58 ± 4.25%, n=4; *Gdnf*^wt/hyper^;*Wnt11*^+/+^: 75.45 ± 2.44%, n=12; *Gdnf*^wt/hyper^;*Wnt11*^+/−^: 73.52 ± 2.74%, n=19; *Gdnf*^wt/hyper^;*Wnt11*^−/−^: 56.52 ± 7.47%, n=2; *Gdnf*^hyper/hyper^;*Wnt11*^+/+^: 46.90 ± 4.16%, n=8; *Gdnf*^hyper/hyper^;*Wnt11*^+/−^: 41.67 ± 3.06%, n=7;) The estimated marginal means of kidney size in (A) and (B) are results of the analysis containing all the data about kidney size in (C). Kidney size in *Gdnf*^wt/hyper^*;Wnt11*^−/−^ is significantly smaller than that in *Gdnf*^wt/hyper^*;Wnt11*^wt/wt^ mutants (P=0.014, unpaired 2-tailed t-test). (D) Representative H&E staining image showing morphology in cortical region of *Gdnf*^hyper/hyper^;*Wnt11*^+/+^ and (E) *Gdnf*^hyper/hyper^;*Wnt11*^+/−^ kidney at P0. The dilated UB (red arrow), collecting duct cysts (black arrow head), cyst in glomerular tuft (black arrow) are slightly improved in *Gdnf*^hyper/hyper^;*Wnt11*^+/−^ kidneys than those seen in *Gdnf*^hyper/hyper^. * denotes P < 0.05, ** denotes P < 0.01, *** denotes P < 0.001 in comparison with each other; one-way ANOVA followed by Bonferroni’s *post-hoc* test. *Scale bar*: 100μm.

**Table S1.** The primary and secondary antibodies used in this study.

| **Primary Antibody** | **Host** | **Dilution** | **Manufacturer** |
| --- | --- | --- | --- |
| CALBINDIN | goat pAb | 1:500 | Santa Cruz Biotechnology |
| CYCLIN D1 | rabbit mAb | 1:400 | NeoMarkers (Thermo Scientific) |
| EPITHELIAL CADHERIN | goat pAb | 1:500 | R&D Systems |
| JAG1 | rat mAb | 1:200 | Hybridoma Bank |
| Ki67 | rabbit mAb | 1:200 | Abcam |
| LEF1 | rabbit mAb | 1:400 | Cell Signaling Technology |
| PAX2 | rabbit pAb | 1:400 | Invitrogen |
| PODOCALYXIN | goat pAb | 1:300 | R&D Systems |
| SIX2 | rabbit pAb | 1:500 | ProteinTech |
| SOX9 | rabbit pAb | 1:500 | Millipore |

| *Secondary Antibody* | *Host* | *Dilution* | *Manufacturer* |
| --- | --- | --- | --- |
| anti-goat Alexa Fluor® 488 | Donkey | 1:400 | Jackson Immuno Research Laboratories |
| anti-rat Cy™3 | Donkey | 1:400 | Jackson Immuno Research Laboratories |
| anti-rabbit Alexa Fluor® 568 | Donkey | 1:400 | Jackson Immuno Research Laboratories |
| anti-goat Alexa Fluor® 647 | Donkey | 1:400 | Jackson Immuno Research Laboratories |
| anti-rabbit IgG (H+L) - Horseradish Peroxidase (HRP) conjugate | Goat | 1:2000 | Bio-Rad |
| anti-goat IgG (H+L) - Horseradish Peroxidase (HRP) conjugate | Rabbit | 1:500 | Bio-Rad |

**Table S2.**
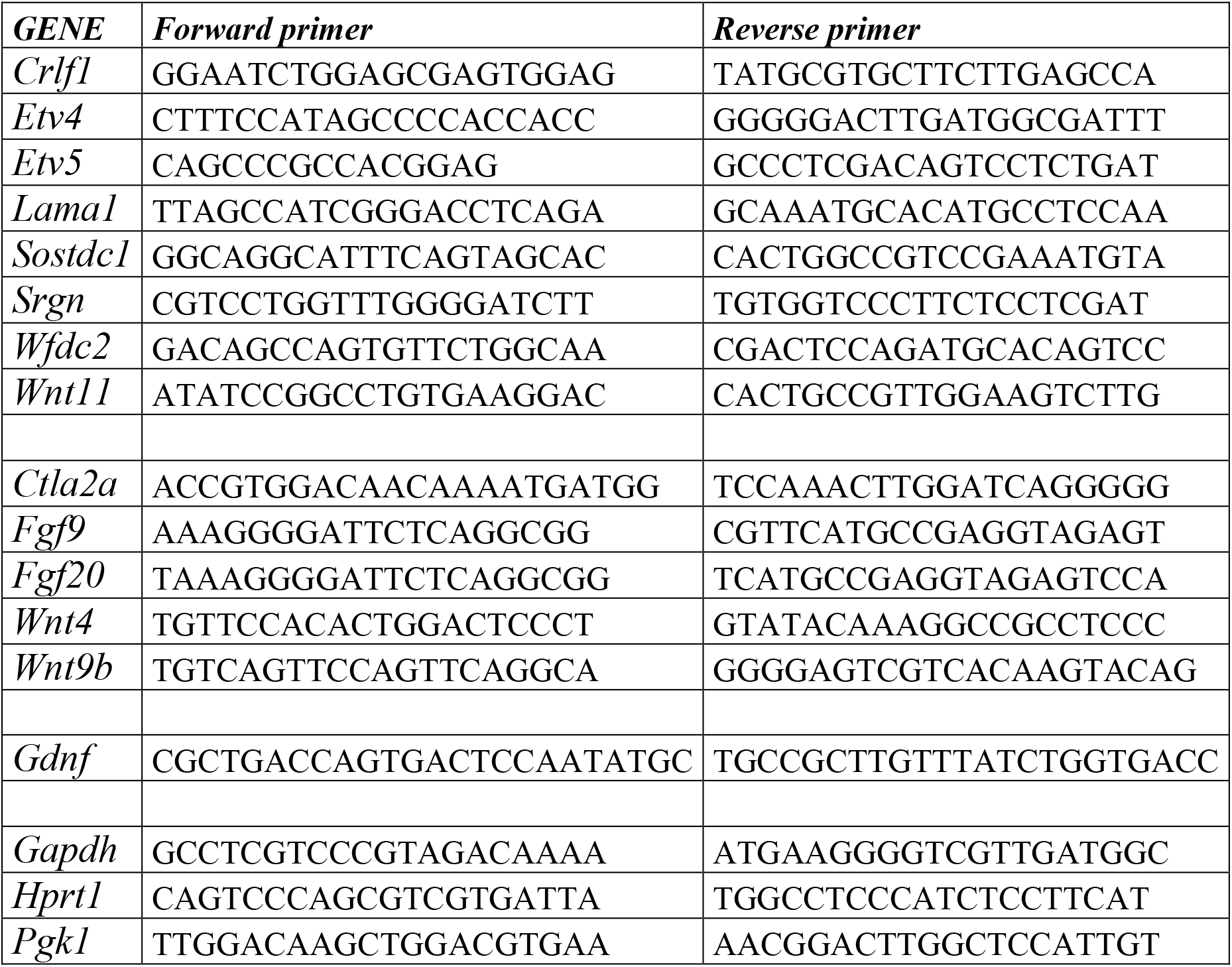
The primers used for qRT-PCR in this study.

